# Polymerase Eta Recruits FANCD2 to Common Fragile Sites to Maintain Genome Stability

**DOI:** 10.1101/2025.01.06.631600

**Authors:** Mrunmai Niljikar, Angelica Barreto-Galvez, Saloni Patel, Julia Elizabeth Gagliardi, Vasudha Kumar, Archana Pradeep, Aastha Juwarwala, Jeannine Gerhardt, Yokechen Chang, Cristina Montagna, Advaitha Madireddy

**Affiliations:** Rutgers Cancer Institute of New Jersey, New Brunswick, NJ, USA; Department of Obstetrics and Gynecology, Weill Cornell Medicine, New York, NY, USA; Department of Genetics, Albert Einstein College of Medicine, Bronx, NY, USA; Department of Pediatrics Hematology/Oncology, Robert Wood Johnson Medical School, Rutgers University, New Brunswick, NJ, USA

## Abstract

The replicative polymerase delta is inefficient copying repetitive DNA sequences. Error-prone translesion polymerases have been shown to switch with high-fidelity replicative polymerases to help navigate repetitive DNA. We and others have demonstrated the importance of one such translesion polymerase, polymerase Eta (pol eta), in facilitating replication at genomic regions called common fragile sites (CFS), which are difficult-to-replicate genomic regions that are hypersensitive to replication stress. However, the mechanistic basis for pol eta’s role in facilitating DNA replication at CFS and(or) at other genomic regions is currently unclear. Importantly, the functional importance of three non-catalytic domains of pol eta, the Ubiquitin-binding Zinc finger (UBZ), PCNA interacting protein (PIP) domain, and the F1 domain which mediates its switch with replicative DNA polymerases in mediating replication stress, especially at CFS loci is not clear. Here, we report that the PIP and UBZ domains of Pol Eta are both critical for its role in mediating cellular replication stress, especially at CFS. The absence of either domain induced elevated replication stress, replication stalling and DNA damage accumulation genome wide. This effect was even more pronounced at CFS loci leading to the accumulation of under replication DNA in G2/M. Importantly, while the inactivation of the UBZ domain resulted in a robust FANCD2 monoubiquitylation (a prominent marker of FANCD2 activation), FANCD2 recruitment genome wide was significantly impacted, especially at CFSs such as FRA16D. These S-phase phenotypes result in ssDNA gap formation and the persistence of under-replicated genomic regions upon transition to G2/M. While post-replicative gap filing/ repair by Mitotic DNA synthesis is activated in the mutants, it only effectively resolves UFBs in the F1* cells. The PIP*, UBZ* and pol eta^−/−^ cells unfortunately manifest excessive toxic cytosolic DNA that instigates a strong innate immune response. These results collectively show that translesion polymerase Eta functions in a common pathway with FANCD2 to prevent replication perturbation and instability at CFS loci.

## Introduction

Nearly two-thirds of the human genome is comprised of repetitive DNA elements^1^, which inherently challenge the movement of replication machinery, threatening genome integrity. Common fragile sites (CFS) are genomic regions that contain repetitive DNA that are hypermutable in response to replicative stress^2^. We have previously shown that CFS displays spontaneous chromosomal aberrations in response to endogenous stress^3^. Particularly, we and others have demonstrated that CFS fragility is a cell-type specific phenomenon, attributed to perturbed DNA replication and gene transcription^4,5^, and strongly implicated in carcinogenesis^6,7^. In fact, hypermutations and chromosomal abnormalities at CFS, such as deletions and translocations are prevalent in a number of cancers^7–9^.

The inherent vulnerability conferred by replication challenges found at CFSs suggests the existence of a protective cellular network that prevents instability even under normal replicative conditions. Indeed, studies from our lab and others have revealed that fragile site replication is perturbed significantly, leading to spontaneous fragility at these regions in the absence of a functional Fanconi anemia (FA)/BRCA pathway^3,10,11^ in human cells. Importantly, while we discovered FANCD2 to be an important mediator of CFS stability, the signaling networks surrounding FANCD2’s recruitment and functional role to enable CFS replication completion to prevent instability is still unclear. Replicative polymerases by nature are inadept at replicating the repetitive DNA that are present at CFS regions. A specialized class of polymerases involved in translesion synthesis (TLS), have been postulated to switch with high fidelity replicative polymerases, to maintain CFS stability and prevent chromosomal breakages at CFSs^12–14^. Importantly, we and others have recently shown that Translesion polymerase Eta (Pol Eta) deficiency results in replication stalling and increased dormant origin activation at CFS regions, making it a critical component of the replication machinery at CFS^15,16^. However, several questions regarding its mechanism of action remain unanswered.

For instance, it is not clear whether the non-catalytic domains of Pol Eta, such as the Ubiquitin-binding Zinc finger (UBZ), PCNA interacting protein (PIP) domain, and the F1 domain which mediates its switch with replicative DNA polymerases are required for its CFS function. Furthermore, it is not known whether Pol Eta functions by itself or in consort with other proteins such as FANCD2 to ensure CFS stability. Here, we report that the PIP and UBZ domains of Pol Eta are both critical for its role in mediating cellular replication stress, especially at CFS. The absence of either domain induced elevated replication stress, replication stalling and DNA damage accumulation genome wide. This effect was even more pronounced at CFS loci leading to the accumulation of under replication DNA in G2/M. Importantly, while the inactivation of the UBZ domain resulted in a robust FANCD2 monoubiquitylation (a prominent marker of FANCD2 activation), FANCD2 recruitment genome wide was significantly impacted, especially at CFSs such as FRA16D. These S-phase phenotypes result in ssDNA gap formation and the persistence of under-replicated genomic regions upon transition to G2/M. While post-replicative gap filing/ repair by Mitotic DNA synthesis is activated in the mutants, it only effectively resolves UFBs in the F1* cells. The PIP*, UBZ* and pol eta^−/−^ cells unfortunately manifest excessive toxic cytosolic DNA that instigates a strong innate immune response. These results collectively show that translesion polymerase Eta functions in a common pathway with FANCD2 to prevent replication perturbation and instability at CFS loci.

## Results

### Polη deficiency closely resembles FANCD2 deficiency at CFS

Previous research from our lab showed perturbed replication dynamics at the CFS-FRA16D locus in the absence of the FANCD2 protein in lymphocytes3. Since CFS are cell-type specific, we first established replication dynamics at a bonafide fibroblast specific fragile sites called NFIA (Fig. 1A) in non-affected IMR90 fibroblasts. Replication at the NFIA locus in the IMR90 fibroblasts was largely carried out by forks moving in the 3’ to 5’ direction with several terminating forks. As expected, no initiation events were observed in this fragile site region (Fig. 1B). In contrast, FANCD2 deficient fibroblasts replicated the NFIA regions with forks moving predominantly (66%) in the 5’ to 3’ direction (Fig. 1C). In addition, several initiation events were observed in the absence of FANCD2. Analysis of replication dynamics in a Polη deficient fibroblast (Fig. 1D) at the same locus revealed very similar replication dynamics to FANCD2 deficient cells with a majority of forks moving in the 5’ to 3’ direction (Fig. 1E). The similar perturbations in replications dynamics at NFIA in the absence of either protein is suggestive of a common pathway involving the two to ensure replication integrity at CFS loci.

**Figure 1:**
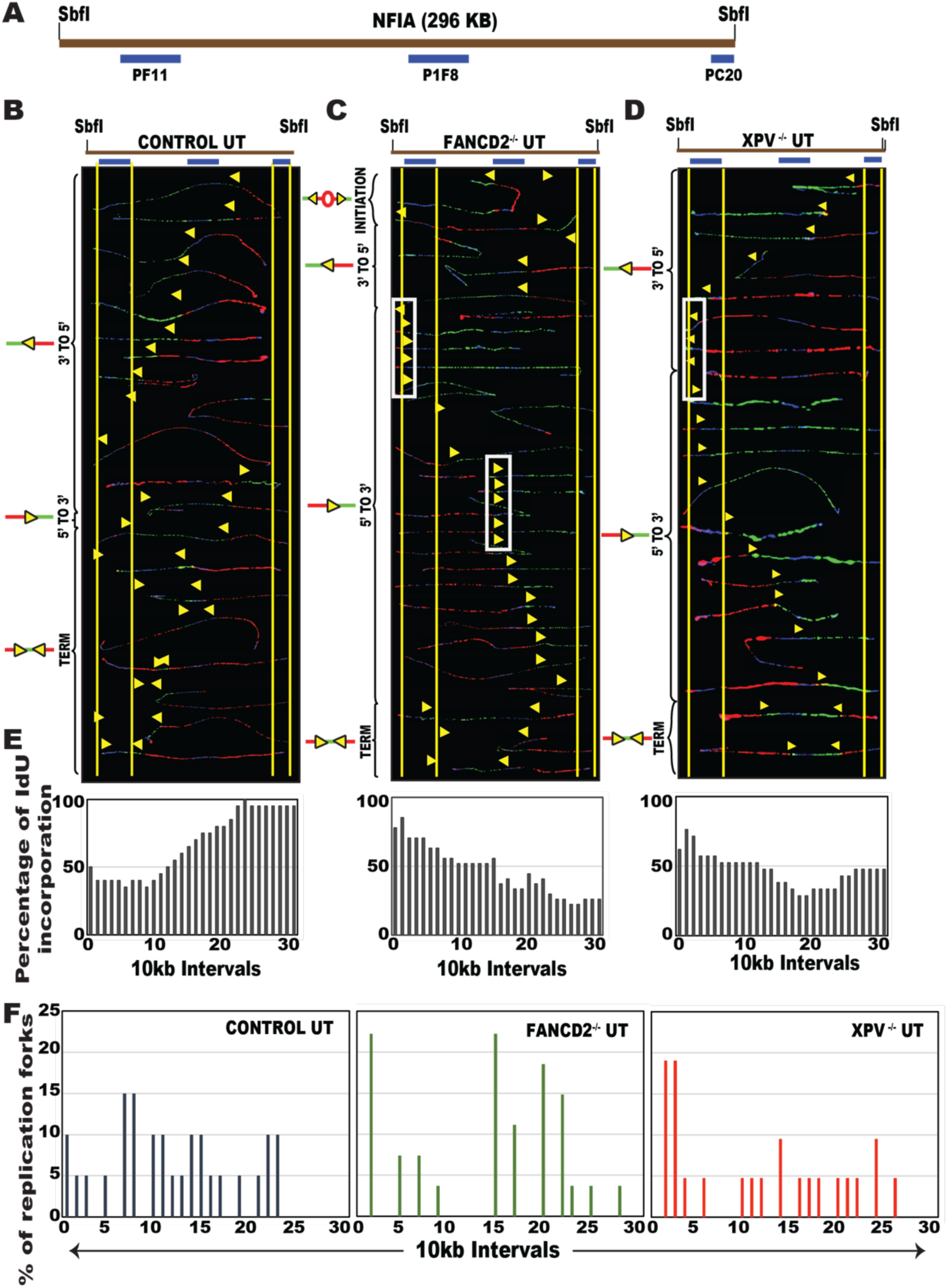
Polη deficiency closely resembles FANCD2 deficiency at CFS. **Top:** (A) Locus map of a 296 kb region in the CFS-NFIA obtained by SbfI digestion. The FISH probes that identify the segment are labeled in blue. Top; Locus map of the SbfI digested NFIA segment. Bottom; Aligned photomicrograph images of labeled DNA molecules from DNA Replication program at CFS-NFIA in IMR90 fibroblasts (B), FANCD2^−/−^ fibroblasts(C) and XPV^−/−^ fibroblasts(D). The yellow arrows indicate the sites along the molecules where the IdU transitioned to CldU. The yellow arrows with white ovals indicate replication fork origin activation. The molecules are arranged in the following order: molecules with initiation events, molecules with 3’-to-5’ progressing forks, molecules with 5’-to-3’ progressing forks, and molecules with termination events. (E) The percentage of molecules incorporating IdU (red) is calculated from the replication program (middle) and is represented as a histogram. (F) The percentage of molecules with replication forks at each 10 kb interval of NFIA. Black arrows denote the most prominent pause peaks and correspond to the white ovals in the SMARD profile.

### The non-catalytic domains of Polη, especially the UBZ domain, are essential for its role facilitating replication across CFS-NFIA

To further understand the mechanistic basis for Pol Etas role in the replication stress response, especially at CFS loci, we evaluated five isogenic fibroblast cell lines were generated from Polη deficient XP30RO patient derived fibroblasts (pol eta^−/−^) using Polη mutant constructs. Polη has various protein domains-the catalytic domain, required for DNA synthesis and three non-catalytic domains (PIP, UBZ and F1) that are implicated in Polη’s recruitment and its interaction with other proteins (Fig. 2A). The UBZ mutant involves a D-to-A substitution at position 652 (D652A), which inhibits Polη’s ability to interact with monoubiquitinated proteins like PCNA and possibly FANCD2^17^. The PIP mutant consists of a deletion of the last nine amino acids generated by introducing a stop codon at position 705 promoting a deletion of the last nine amino acids of the Polη protein. This has been shown to inhibit interaction between Polη and PCNA^12,18^. The F1 mutant involves mutations in the FF_483–484_ amino acid residues of Polη (designated F1 motif). These residues are essential for the interaction of Polη with POLD2, the B subunit of the replicative DNA polymerase δ, both *in vitro* and *in vivo*^19^. It is known that that the F1 motif is required for the progression of DNA synthesis after UV irradiation and efficient TLS complex^19^. To ensure stable and robust Polη protein expression in all five isogenic cell lines, we first carried out an immunoblot analysis. As expected, the pol eta^−/−^ cells were completely deficient for Polη protein expression (Fig. 2B), all three mutants and the functionally complemented wildtype (Pol eta^+/+^) display Polη expression.

**Figure 2:**
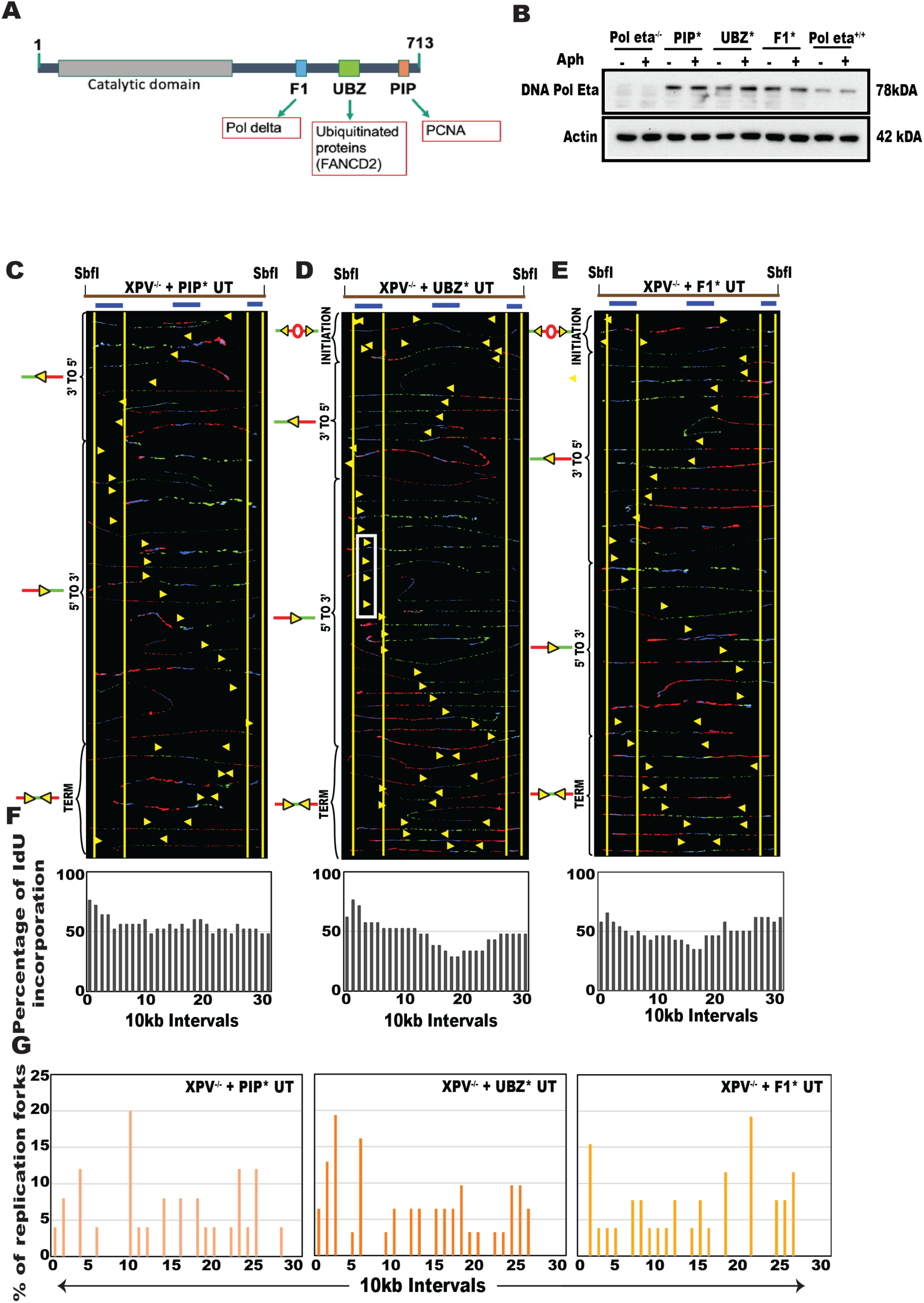
The non-catalytic domains of Polη, especially the UBZ domain, are essential for its role facilitating replication across CFS-NFIA. **Top: Pol eta structure and expression in** (A) Protein map of Structure of DNA Polymerase eta with its protein domains (Catalytic, UBZ, PIP and F1). (B) Expression levels of Polη and Actin proteins in isogenic cell line models of Polη treated with APH by detected by western blotting. **Middle**: Aligned photomicrograph images of labeled DNA molecules from DNA Replication program at CFS-NFIA in (C) XPV^−/−^ +PIP* fibroblasts; (D) XPV^−/−^ +UBZ* fibroblasts; and (E) XPV^−/−^ +F1* fibroblasts. The yellow arrows indicate the sites along the molecules where the IdU transitioned to CldU. The molecules are arranged in the following order: molecules with initiation events, molecules with 3’ to 5’ travelling forks, molecules with 5’ to 3’ travelling forks and molecules with termination events; **Bottom:** Quantifications for (F) Percentage of molecules with IdU incorporation and (G) Percentage of replication forks in XPV^−/−^ +PIP* fibroblasts, XPV^−/−^ +UBZ* fibroblasts and XPV^−/−^ +F1* fibroblasts.

To assess the functional impact each non-catalytic domain of Pol Eta on replication facilitation through CFS loci, we assessed the replication program of NFIA in these isogenic cell lines expressing the PIP, UBZ and F1 mutated Pol Eta (Fig. 2C-G). The results showed that the NFIA locus in both the PIP and UBZ domain mutants was replicated predominantly by forks moving in the 5’ to 3’ direction (Fig. 2C-D), very similar to the FANCD2 deficient line (Fig. 1C). In contrast, the F1 domain mutant appeared to have relatively normal replication dynamics being replicated by equal numbers of forks travelling in either direction (Fig. 2E). However, there was a notable increase in replication initiation events in the UBZ* and F1* cells, indicative of elevated basal replication stress even under unperturbed conditions. The percentage of terminating forks although high, were quite similar between the three pol eta mutants. While replication across CFS-NFIA appears to be perturbed to a similar degree in cells deficient for Pol Eta or expressing mutant Pol Eta, the UBZ* cells that had prominent replication pausing and dormant origin activation, displayed the closest resemblance to FANCD2 deficient cells. Overall, these results suggest that the UBZ and F1 domains of Pol Eta are important for its role at CFS loci.

### Loss of the non-catalytic domains of Polη but not Polη itself enhances the sensitivity of cells to replication stress

Replicative difficulties during the S-phase result in activation of ATR-Chk1 signaling cascade^20^. To understand whether the replication perturbation observed in the mutants at CFS-NFIA extends genome wide, we first carried out an immunoblot analysis to assess spontaneous and induced ATR/Chk1 phosphorylation. Aphidicolin (APH), a polymerase alpha inhibitor was used in low doses to induce replication stress. The analysis revealed a spontaneous phosphorylation of ATR and Chk1 in all cell lines, including the WT Pol eta complemented cells (Pol eta^+/+^). Upon exposure to replication stress, this response was further exacerbated in all mutants, with the exception of Pol eta−/−, which had similar band intensity with and without inhibitor treatment. Interestingly, loss of the catalytic domains of pol eta, especially the UBZ* and F1*, resulted in the strongest spontaneous and induced pChk1 activation, even more so than cells completely lacking the pol eta protein. To determine whether checkpoint activation is a result of replication fork-stalling genome wide, we carried out a DNA fiber analysis^21^. Here, cells were pulse labelled with nucleoside analog IdU for 30 mins followed by CldU for 60 mins during which a replicative inhibitor is added to assess fork stalling. A shortening of the CldU tract (indicated by a ratio <3) would be indicative of replication stalling. Results from this analysis showed that HU treatment resulted in replication fork stalling in all the cell lines analyzed. As expected, the wildtype Pol eta complemented cells showed the least amount of fork stalling upon HU treatment. While all non-catalytic domain mutants displayed significantly shorter CldU tracts as compared to pol eta wildtype cells, a comparison of the extent of stalling between the treated pol eta mutant pairs revealed significantly shorter CldU lengths in UBZ* and PIP* cells compared to the pol eta−/− and F1* cells. These results suggest that the UBZ and PIP domains of pol eta are important in alleviating replication fork pausing upon replication stress.

### PIP* and UBZ* cells have elevated persistent unresolved DNA damage

Next, to determine whether increased replication pausing resulted in elevated genomic instability in these cells, we measured the overall magnitude of spontaneous and induced genomic instability by immunostaining (IF) for phosphorylation of histone variant 2A (pH2AX). As expected, the complete absence of Pol eta resulted in a modest but significant increase in DNA damage in response to replication stress as compared to cells with intact wildtype Pol Eta (**Fig. 3C**; compare red to green). However, the presence of mutant pol eta, especially the PIP and UBZ domain mutated pol eta, was far more deleterious to the cells even in the absence of exogenous replication stress. This effect was only further exacerbated upon replicative inhibition (**Fig. 3C**). We then validated instability at the single cell level by measuring DNA breaks using the Comet assay^22^. Similar to the IF results, we observed a significant increase in spontaneous and stress induced comet tail movement in pol eta mutated cells (orange), compared to the pol eta wildtype cells (**Fig. 3D**).

**Figure 3:**
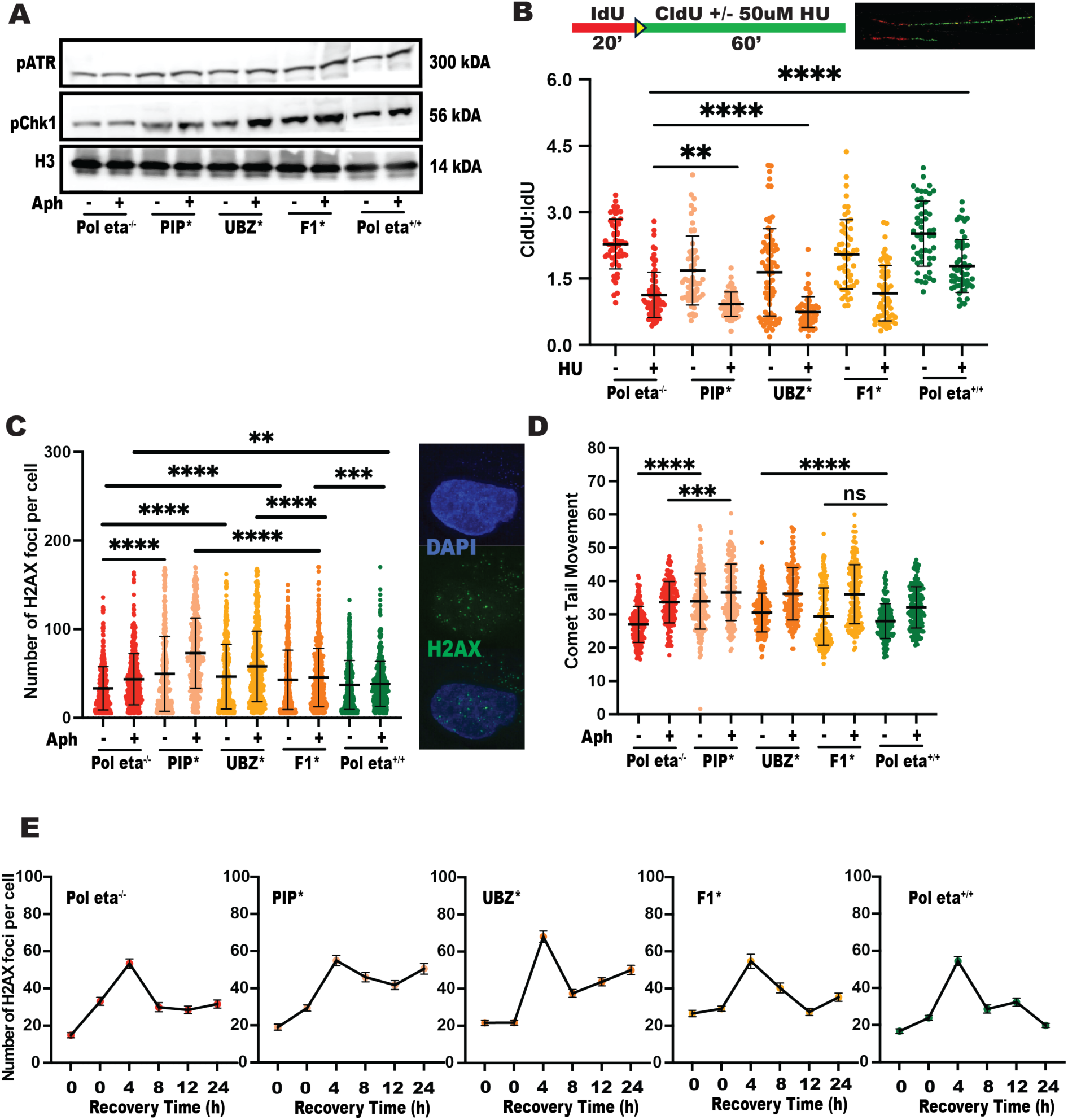
Loss of the non-catalytic domains of Polη but not Polη itself enhances the sensitivity of cells to replication stress. (A) Measuring expression levels of phospho-ATR protein, phospho-Chk1 Ser317 to assess replication checkpoint activation using western blotting. Expression levels of H3 were used as a loading control and (B) DNA fiber analysis of 50uM hydroxyurea (HU) treated to assess replication fork stalling in pol eta^−/−^, PIP*, UBZ*, F1* and pol eta+^/+^ cells. The fork rate (CldU/IdU ratio) is indicated, n=150. The p-values are indicated as follows: * <0.03, ** <0.0021, *** <0.0002, **** <0.0001. Scale bar 10 μm; (C) Analysis of number of gH2AX foci (red) in the nucleus per cell in Pol eta^−/−^, PIP*, UBZ*, F1* and Pol eta^+/+^ cells when exposed to Aphidicolin (Aph). Representative images are on the top, n=250; The p-values are indicated as follows: * <0.03, ** <0.0021, *** <0.0002, **** <0.0001. (D) Measurement of DNA single strand breaks by alkaline Comet assay in Pol eta^−/−^, PIP*, UBZ*, F1* and Pol eta^+/+^ cells treated with Aphidicolin (APH). The p-values are indicated as follows: * <0.03, ** <0.0021, *** <0.0002, **** <0.0001. Lengths were measured using the Adobe Photoshop Tools. (E) Time course experiment to measure recovery of cells after release into drug free media over the course of 48hours. Cells were collected at six time points (0, 4, 8, 12, 24, 28 hours) and expression levels of phospho-Histone H2AX Ser139 were measured by Immunofluorescence staining. The p-values are indicated as follows: * <0.03, ** <0.0021, *** <0.0002, **** <0.0001. Scale bar 10 μm.

Next, to assess the proficiency of DNA damage resolution in the pol eta mutants, we carried out a time course experiment. In short, the entire isogenic panel of cell lines were treated with low does APH for 16 hours followed by which the cells were released into drug free media for recovery. The cells were then collected at different timepoints (0, 4, 8, 12, 24h) and subjected to IF and immunoblotting to assess both damage and checkpoint resolution. The results from this analysis showed a clear separation in damage resolution capabilities between the pol eta non-catalytic domain mutants as compared to pol eta proficient and even pol eta deficient cells (**Fig. 3E**). While the average basal level of pH2AX foci per cell was less than 20 in pol eta^−/−^ and pol eta^+/+^ cells, all three mutants has ~20-30 spontaneous pH2AX foci per cell. Upon replicative inhibition, the slight increase in pH2AX foci observed in all cell lines peaked to between 55-60 foci per cell in the pol eta^−/−^, pol eta^+/+^, PIP* and F1 cells. The UBZ* cells accumulated a significantly higher number of pH2AX (~70+) foci per cell at the 4h timepoint as compared at all other cells. In addition, the pH2AX foci number started decreasing at the 8h time point in all cell lines but only efficiently resolved by 24h in the pol eta^+/+^ wildtype cells (**Fig. 3E**). The pol eta^−/−^ and F* cells displayed 50% decrease in the number of H2AX foci by 24h (~30 foci), the damage did not completely resolve at the point. This effect was further exacerbated in the PIP* and UBZ* cells which only showed marginal decrease in foci per cell which persisted even at the 24h timepoint. These studies indicate that the UBZ domain and to a lesser extent the PIP domain is both important for either DNA damage tolerance or repair in response to replication stress.

### Cells harboring inactivating mutations in the non-catalytic domains of Polη prefer repriming over translesion synthesis for DNA damage tolerance

While cells employ several well characterized DNA repair mechanisms to remove a lesion that stalls replication, the quick bypass of the lesion or impediment to replicative polymerases are carried out by DNA damage tolerance (DDT) pathways such as translesion synthesis (TLS), Template switching or repriming^23^. While DDT only allows lesion bypass, processes such as TLS reduce instances of replisome pausing, thus reducing the chances of fork collapse. Under basal levels of stress (metabolic-endogenous), stalling of replication forks can occur at secondary structure prone repetitive DNA that can fold into non-B DNA structures. These structures while difficult to replicate by replicative DNA polymerases, are easily handled by TLS polymerases that can accommodate large lesions within their active sites. Considering the inherently higher levels of endogenous stress, replisome pausing and damage accumulation in pol eta^−/−^, PIP*, UBZ* and F1* cells, we assessed the reliance of these cells on TLS as the primary mechanism of DDT. Ubiquitinated proliferating cell nuclear antigen (ubPCNA), is considered a direct measure of TLS activity as it signals the recruitment of TLS polymerases to the site of DNA damage, allowing for replication to continue past lesions. To assess TLS activity, we carried out an immunoblot analysis to measure the extent of PCNA ubiquitination. While pol eta^−/−^ and pol eta^+/+^ cells showed a robust spontaneous and stress-induced PCNA ubiquitination, it was very interesting to note that all three Pol eta domain mutants displayed a very low level of PCNA ubiquitination (**Fig. 4A**). Importantly, the amount of ubPCNA was inversely proportional to the amount of replication stalling and DNA damage observed. This phenotype was not a factor of altered PCNA levels since the overall levels of PCNA protein expression were consistently the same between all the isogenic cell lines evaluated. Next, we evaluated pol eta chromatin recruitment as a second measure to confirm the degree of cellular reliance on TLS in each cell line. As expected, pol eta chromatin recruitment was absent in the pol eta^−/−^ cells and robust pol eta foci formation was observed in pol eta^+/+^ cells both spontaneously and in the presence of replication stress (**Fig. 4B**). In the three mutants, the number of pol eta foci observed mimicked the extent of ubPCNA detected. For example, while the PIP* cells had a considerably lower number of pol eta foci under both untreated and treated conditions, treated UBZ* cells had a significant increase in pol eta foci (**Fig. 4B**) which was represented by the prominent ubPCNA band detected (**Fig. 4A**). These results suggest that the extent of replication stress likely determines the DNA damage tolerance/repair activated by the cells to avoid fork collapse. They also indicate that an alternate DDT mechanism or stalled fork restart mechanism is utilized by cells harboring mutations in the non-catalytic domains of Polη.

**Figure 4:**
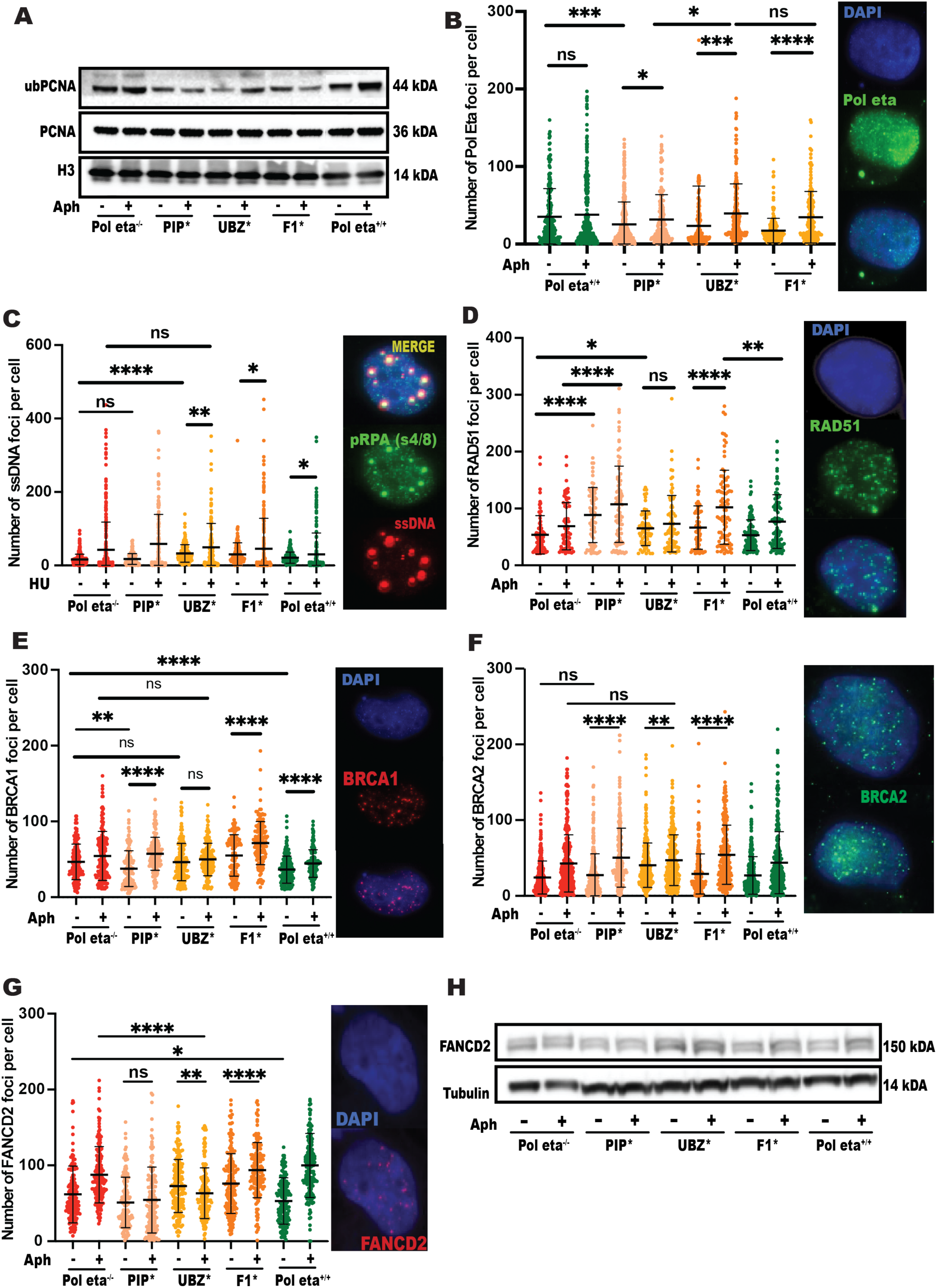
PIP* and F1* but not UBZ* rely on FA/BRCA mediated fork protection in response to replication stress. (A) Expression levels of ubPCNA and PCNA in Pol eta^−/−^, PIP*, UBZ*, F1* and Pol eta^+/+^ cells treated with and without Aphidicolin (APH) using western blotting. H3 is used as loading control. (B) Expression levels of Pol eta in PIP*, UBZ*, F1* and Pol eta^+/+^ cells treated with and without Aphidicolin (APH) using immunofluorescence staining. (C-F) Chromatin recruitment of RAD51, BRCA1, BRCA2 and FANCD2 proteins assessed using immunofluorescence staining. Representative images are depicted on the right. (G) FANCD2 mono-ubiquitination determined by western blotting in Pol eta^−/−^, PIP*, UBZ*, F1* and Pol eta^+/+^ cells treated with Aphidicolin (APH).

Pol eta mutant cells (PIP*, UBZ* and F1*) exhibit elevated replication pausing as compared to both pol eta^−/−^ and pol eta^+/+^ cells, which in turn was associated with increased unresolved DNA damage. However, mutant cells did not activate TLS for DNA damage tolerance (DDT). An alternate mechanism of DDT involves repriming by PRIMPOL^24–27^. Repriming involves re-initiation of DNA synthesis beyond a DNA lesion, leaving unreplicated single-stranded DNA (ssDNA) gaps to be filled post-replicatively^28^. To determine whether the pol eta mutant cells (PIP*, UBZ* and F1*) rely on repriming to tolerate DNA damage, we assessed ssDNA accumulation, a common byproduct of PRIMPOL activity, by IF. Under unperturbed conditions, the UBZ* and F1* mutants showed a significant upregulation in ssDNA gap accumulation as compared to PIP*, pol eta^−/−^ and pol eta^+/+^ cells (**Fig. 4C**). Upon exposure to replication stress (hydroxyurea – HU treatment), pol eta^−/−^, PIP* and UBZ* showed a significant increase in ssDNA gaps as compared to F1* and pol eta^+/+^ cells, indicating a potential upregulation of repriming for DDT.

### PIP* and F1* but not UBZ* rely on FA/BRCA mediated fork protection in response to replication stress

In normal cells, replication fork stalling in response to replication inhibition is most often handled by the FA/BRCA fork protection pathway^29^. This error-free process involves fork reversal at nascently synthesized DNA followed by RAD51 loading and the efficient restart of stalled forks in the presence of downstream effectors such as BRCA1/BRCA2/ PALB2, thus maintaining genome stability. Next, we evaluated whether pol eta non-catalytic domain mutants utilized fork protection as a means of restarting stalled replication. First, we assessed chromatin recruitment of RAD51, BRCA1 and BRCA2 by immunofluorescence staining. The PIP* and F1* cells, which showed the lowest levels of ubPCNA displayed a significant increase in chromatin recruitment of RAD51, BRCA1 and BRCA2 under both treated and untreated conditions (**Fig. 4D-F**; peach and orange). While the untreated UBZ* cells which had a low basal level of ubPCNA expression had an inherently higher chromatin recruitment of RAD51, BRCA1 and BRCA2, as compared to the pol eta^−/−^ and pol eta^+/+^ cells, there was no significant increase in RAD51 or BRCA1 and a modest increase in BRCA2 foci formation in response to replication stress (**Fig. 4D-F**; yellow). Taken together, these results suggest that UBZ* cells rely on fork protection at basal levels but are likely employing an alternate mechanism for DDT or fork restart upon replication stress.

### Inactivation of the UBZ domain of Polη results in decreased recruitment of FANCD2 to the chromatin

Another important component of the FA/BRCA fork restart machinery is a FA protein called FANCD2, which^29^. Of note, it has been shown that FANCD2 nuclear foci localize predominantly to^29^ likely due to its role in facilitating efficient replication^3^. Here, to confirm the reliance (or lack thereof) of pol eta mutants on the FA/BRCA fork protection pathway, we measured the efficiency of FANCD2 recruitment to sites of replication stalling by immunofluorescence. The results revealed an increased accumulation of FANCD2 nuclear foci during the S-phase in the presence of aphidicolin in the PIP*, F1*, pol eta^−/−^ and pol eta^+/+^ cells, similar to the RAD51, BRCA1 and BRCA2 foci formation observed previously. The highest FANCD2 foci formation was observed in the absence of F1 domain of Polη likely due to its role in facilitating the switch between replicative DNA polymerases and ^19,30,19,25^. Interestingly, consistent with the RAD51 and BRCA1 results, the UBZ* mutant showed a significant decrease in FANCD2 foci formation after aphidicolin treatment (**Fig. 4G**). In addition to pointing to a decreased reliance on fork protection as a mechanism of stalled fork restart, these results suggest a potential defect in FANCD2 chromatin recruitment in UBZ* cells.

### Decreased FANCD2 chromatin recruitment is not due to a defect in FANCD2 monoubiquitylation

The monoubiquitination of FANCD2-FANCI is a pivotal step essential to activate fork protection and(or) DNA repair during the S-phase^31^. Importantly, mono-ubiquitinated FANCD2 has been previously shown to interact with the UBZ domain of Polη in response to UV damage^32^. While this interaction has not been previously established as a requirement for FANCD2 recruitment in the presence of replication stress, it could explain the decreased recruitment of FANCD2 to chromatin in the UBZ* cells. To rule out that decreased chromatin recruitment of FANCD2 in the UBZ* cell lines was not due to a defect in FANCD2 mono-ubiquitination, we next measured mono-ubiquitination of FANCD2 by immunoblotting analysis. The results showed robust FANCD2 mono-ubiquitination, indicated by a clear double band, where the upper band of slightly higher molecular weight denoted FANCD2 mono-ubiquitination (m-FANCD2) (**Fig. 4H**), in both APH treated and untreated cells. It was very important to note that the strongest spontaneous and stress induced m-FANCD2 bands were observed in the UBZ* cells, a direct measure of elevated stress experienced by the cells expressing UBZ* pol eta. These results clearly show that decreased FANCD2 chromatin recruitment accompanying the overall decrease in chromatin recruitment of other fork protection proteins after APH treatment is not due to a defect in FANCD2 monoubiquitylation. The FANCD2 monoubiquitylation band is a response to replication stress and(or) DNA damage, and it disappears upon efficient damage resolution. To evaluate this, we conducted a time course experiment to measure disappearance of the monoubiquitylation band by immunoblotting and immunofluorescence staining. The results clearly show that while the FANCD2 monoubiquitylation band resolves by 32-48h in the wildtype cells, it persists in the pol eta mutants, with PIP* and UBZ* showing the strongest bands with the longest persistence (**Fig. 5A**). This is a clear indication of damage persistence and delay in resolution in the mutants.

**Figure 5:**
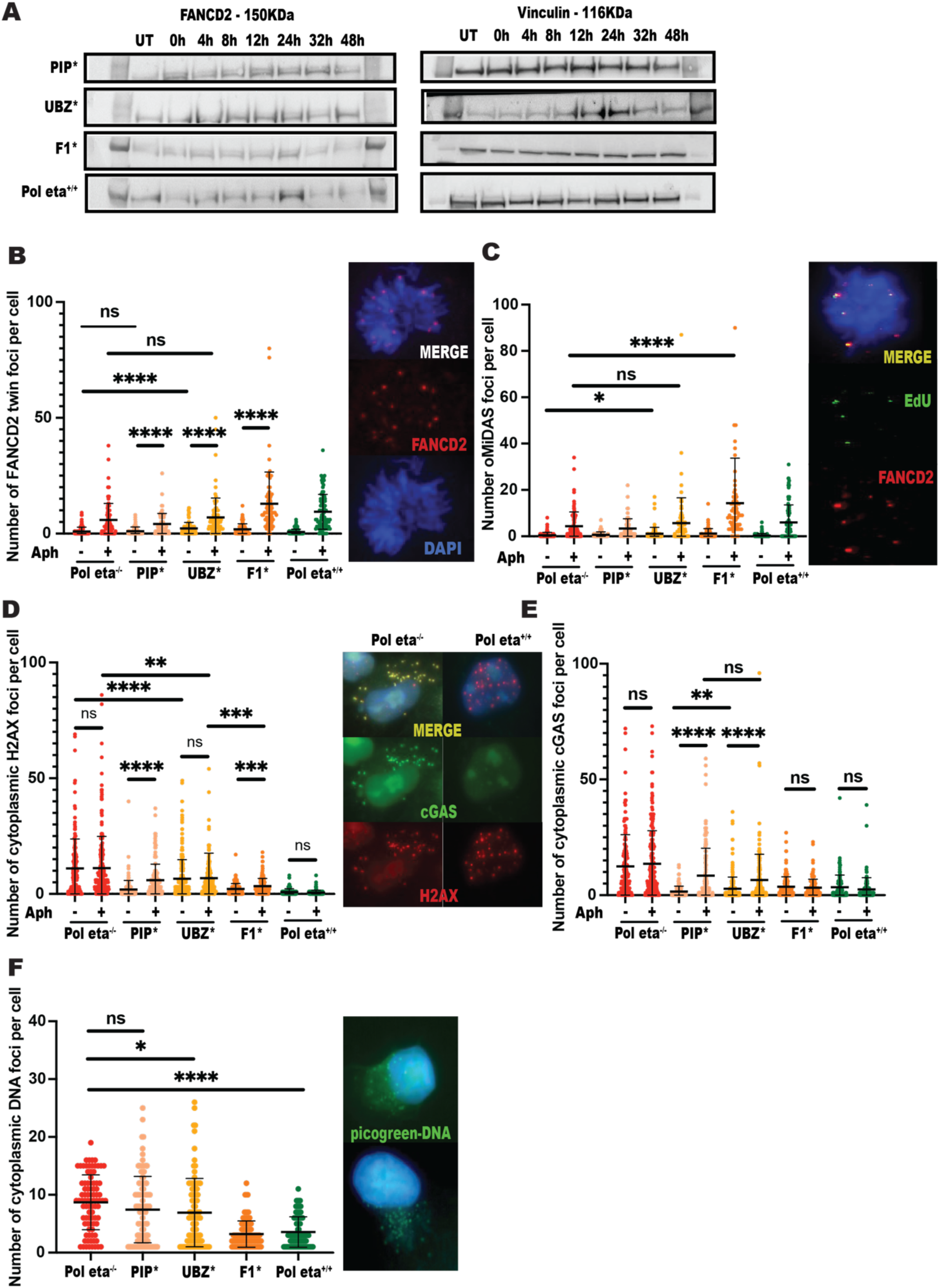
Post-replicative gap filling by Mitotic DNA synthesis and aberrant cytoplasmic DNA accumulation. (A) Time course experiment to measure recovery of cells after release into drug free media over the course of 48 hours. Cells were collected at six time points (0, 4, 8, 12, 24, 28, 36 and 48 hours) and expression levels of FANCD2 mono-ubiquitination was determined by western blotting. (B) Analysis of number of FANCD2 twin-foci (red) per mitotic cell nucleus (DAPI, blue) in Pol−/−, PIP*, UBZ*, F1* and Pol eta+/+ treated with Aphidicolin (APH), representative images are shown on the bottom, n=100; (C) Analysis of number of EdU foci (green), sandwiched between FANCD2 twin-foci(red), per mitotic cell nucleus (DAPI, blue) in Pol−/−, PIP*, UBZ*, F1* and Pol eta+/+ cells treated with Aphidicolin (APH), representative images are shown on the bottom, n=100. (D) Analysis of number of gH2AX foci (red) in the cytoplasm surrounding each cell nucleus in Pol^−/−^, PIP*, UBZ*, F1* and Pol eta^+/+^ cells with or without Aph treatment. Representative images are on the top, n=100; (E) Analysis of number of cGAS (green) in the cytoplasm surrounding each cell nucleus in Pol eta^−/−^, PIP*, UBZ*, F1* and Pol eta^+/+^ cells with or without Aph treatment. Representative images are in the middle, n=100. The p-values are indicated as follows: * <0.03, ** <0.0021, *** <0.0002, **** <0.0001. (F) Analysis of number of Pico green foci in the cytoplasm surrounding each cell nucleus in Pol eta^−/−^, PIP*, UBZ*, F1* and Pol eta^+/+^ cells with or without Aph treatment. Representative images are in the middle, n=100. The p-values are indicated as follows: * <0.03, ** <0.0021, *** <0.0002, **** <0.0001.

### Inactivation of the UBZ and F1 domains elevate the accumulation of under replicated DNA marked by FANCD2 twin foci in G2/M cells

During the S-phase, FANCD2 chromatin recruitment while present was defective in the absence of a functional UBZ domain in Pol Eta. FANCD2 recruited during the S-phase has been implicated primarily in fork protection at stalled replication forks, especially at CFS loci. Next, we wanted to determine whether decreased recruitment of FANCD2 resulted in increased accumulation of under replicated DNA in G2/M cells. Perturbed DNA replication has been associated with incompletely replicated DNA, visualized as ultra-fine DNA bridges (UFB) during mitosis. UFB’s that originate from CFS loci can be identified by twin foci formed by the Fanconi anemia complementation group D2/I (FANCD2/FANCI), proteins at the termini of the UFB on each chromatid^33^. To determine the presence of under-replicated DNA in Polη mutant cells, we measured the persistence of FANCD2 twin foci in mitotic cells. Our results revealed a spontaneous increase in FANCD2 twin-foci in the PIP*, UBZ*, F1* and pol eta^−/−^ cell lines, as compared to the pol eta^+/+^ cells (**Fig. 5B**). Upon replicative inhibition, while complete absence of Polη is associated with a significant increase in FANCD2 twin-foci, the UBZ* and F1* mutants show the most significant increase in FANCD2 twin foci, as compared to all other cell lines. These results indicate that in the absence of functional UBZ and F1 domains, cells are unable to complete replicating DNA, likely at sites of replication stalling leading to the prevalence of under-replicated DNA at CFS in mitosis. Furthermore, it also shows that the UBZ domain of pol eta is dispensable for FANCD2 recruitment to under-replicated DNA in G2/M.

### Post-replicative gap filling by Mitotic DNA synthesis is ineffective in PIP* and UBZ* cells, leading to micronuclei formation

The condensation of incompletely replicated regions triggers the completion of DNA replication during G2/M by the process of Mitotic DNA Synthesis (MiDAS), predominantly at CFS^34,35^. MiDAS requires the coordinated activity of the nucleases, SLX4, RAD52 and the non-catalytic sub-unit of Polymerase delta, POLD3^36–38^. MiDAS is measured by assessing incorporation of EdU at regions marked by FANCD2 twin foci^38,39^. Analysis of MiDAS in the Polη cell lines revealed an increase in localization of EdU synthesis signal sandwiched between FANCD2 twin-foci in all cell lines upon exposure to replicative stress (**Fig. 5C**). Importantly, the inactivation of UBZ and F1 domains of DNA Polη resulted in a significant activation of MiDAS at under-replicated DNA sites, especially in the F1* which harbored the most UFBs. Next, we wanted to determine whether post-replicative gap filling by MiDAS resolved UFBs efficiently in the mutants. Presence of under-replicated DNA in mitosis and chromosome mis-segregation defects in anaphase can lead to micronuclei formation in the subsequent cell cycle^40,41^. To address this, we measured the percentage of cells with micronuclei positively stained for FANCD2, a marker of under-replicated DNA in G2/M. We observed that the PIP* and UBZ* mutants showed a spontaneous increase in FANCD2 +ve micronuclei (data not shown), clearly indicating replication defects as the source of micronuclei formation. Interestingly, the F1* mutant showed the least numbers of FANCD2 positive micronuclei, providing definitive evidence that MiDAS was sufficient to resolve under-replicated genomic regions in these cells in G2/M.

### Aberrant cytoplasmic DNA accumulation and innate immune system activation is observed in PIP*, UBZ* and pol eta^−/−^ cells

Next, to evaluate whether these outcomes result in endogenous cytosolic DNA accumulation in pol eta mutant cells, we assessed the presence of cytosolic DNA called cytoplasmic chromosome fragments (CCF). Endogenous cytosolic DNA are shown to occur in many cancers^40^. Cytosolic DNA of this nature are identified by positive staining for pH2AX and negative staining for 53BP1. The results from this analysis revealed a highly elevated presence of pH2AX (green) in PIP*, UBZ* and pol eta^−/−^ cells (**Fig. 5D**). These results were confirmed by the observed significant accumulation of cytoplasmic dsDNA, as measured by picogreen staining, in PIP*, UBZ* and pol eta^−/−^ cells, as compared to F1* and pol eta^+/+^ cells (**Fig. 5F**). Toxic cytoplasmic DNA is recognized by cGAS (Cyclic GMP–AMP synthase) generating a second messenger that activates the innate immune signaling cGAS-MITA/STING pathway^42^. Cytoplasmic cGAS is often found bound to pH2AX, which is a DNA damage marker ^40,43^. To assess innate immune signaling, we next measured cGAS binding to cytoplasmic pH2AX stained DNA. Similar to the cytoplasmic pH2AX data, the PIP*, UBZ* and pol eta^−/−^ cells showed a significant accumulation of cytoplasmic cGAS foci, as compared to F1* and pol eta^+/+^ cells (**Fig. 5E**).

### The interaction between the UBZ domain of Polη and FANCD2 is essential to prevent under-replicated CFS DNA in G2/M

In non-affected cells, stress induced replication stalling is efficiently resolved leading to CFS stability and over genome stability (**Fig. 6A**). Inactivation of the UBZ domain (and to a certain extent the PIP domain) of Polη results in extensive replication stalling, inefficient fork restart, leading to under-replicated DNA in G2/M. However, while the F1* effectively resolved (**Fig. 6B** - Genomic stability) under-replicated DNA in G2/M by MiDAS, the UBZ and PIP*, like pol eta^−/−^ display persistent damage that leads to mis segregated DNA during mitosis (**Fig. 6B** - Genomic instability). Importantly, The UBZ* mutant like the pol eta^−/−^ cells showed a prominent presence of chromosomal mis-segregation in the form of FANCD2+ micronuclei.

**Figure 6:**
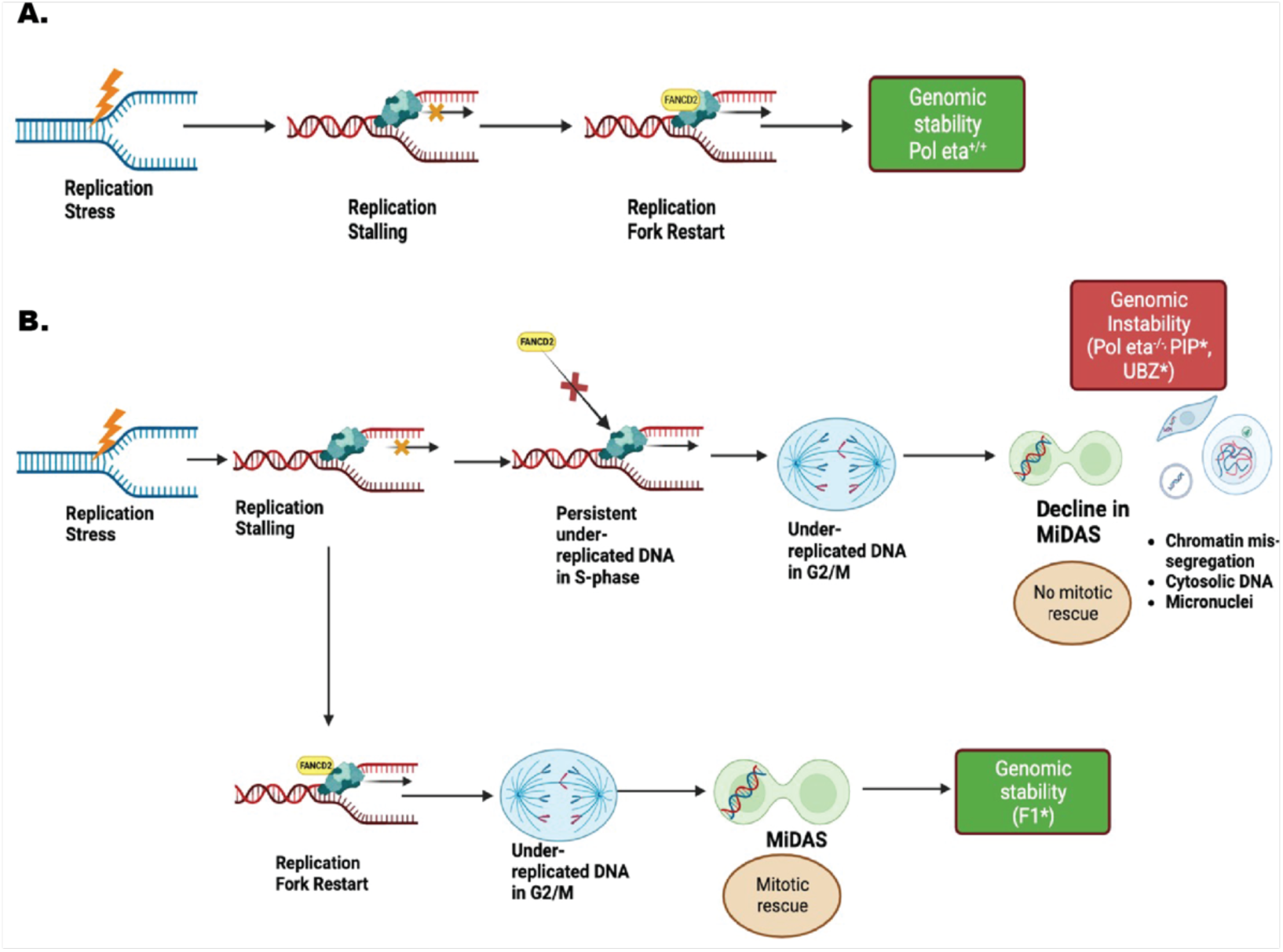
Schematic representation of the working model. (A) In non-affected cells, stress induced replication stalling is efficiently resolved leading to CFS stability and over genome stability. (B) Inactivation of the UBZ domain (and to a certain extent the PIP domain) of Polη results in extensive replication stalling, inefficient fork restart, leading to under-replicated DNA in G2/M. However, while the F1* effectively resolved (Genomic stability) under-replicated DNA in G2/M by MiDAS, the UBZ and PIP*, like pol eta^−/−^ display persistent damage that leads to mis segregated DNA during mitosis (Genomic instability). Importantly, The UBZ* mutant like the pol eta^−/−^ cells showed a prominent presence of chromosomal mis-segregation in the form of FANCD2+ micronuclei.

## Discussion

In our study, we investigated the role of Translesion Polymerase Eta in DNA replication and its importance during replication stress response. Using five isogenic fibroblast cell lines that were generated from a patient derived XPV^−/−^ cell line, we showed that absence of Polη results in altered DNA replication, stalling of the replication fork, enhanced DNA damage, and increased genomic instability. Though the role of Polη has been highly discussed in previous literature of the last decade, the exact mechanism by which it is recruited to sites of DNA damage needs to be explored further. Here, we explored the non-catalytic protein domains of Polη - PIP, UBZ and F1 to evaluate domain-specific roles of the Polη protein in the replication stress response, especially at common fragile sites.

Our results show that loss of the non-catalytic domains of Polη but not Polη itself enhances the sensitivity of cells to replication stress. Although the five human translesion synthesis polymerases have specific sensitivities and lesion specificities, previous literature has shown the functional redundancy between the different translesion polymerases in the absence of one of them. For instance, in circumstances where Polη is completely absent, other TLS polymerases such as pol kappa has been shown to compensate for the deficiency of Polη and to continue replication past an impediment to the replisome. However, in the presence of a mutation in one of the non-catalytical domains of Polη, while it might still get recruited to the site of replisome stalling, it may not be able to resolve the damage, suggesting that having a mutation in one of its protein domains may prove to be more detrimental as compared to its complete absence.

We further found that while the switch between pol delta and eta mediated by the F1 domain is critical to ensure replication completion genome wide, the PIP and UBZ domains of Pol Eta seem critical for recruitment of itself or other proteins to specific genomic regions or perhaps at specific lesions that form more frequently at defined genomic regions. For instance, the CFS loci are prone to specific secondary structures that impede replication fork movement such as non-B DNA structures stemming from repetitive DNA (hairpin, stem loops) and non-B DNA stemming from stalled transcription (DNA: RNA hybrids; R-loops). We have previously shown that FANCD2 is essential to prevent DNA: RNA hybrid formation and(or) stabilization at CFS loci. In addition, we have also previously shown that Pol Eta is essential to maintain stability at CFS loci. While most studies evaluate a functional, mainly catalytic role for pol eta in maintaining CFS stability, our study shows that pol eta might be stabilizing CFS by the mere recruitment of FANCD2 in a non-canonical role. The inactivation of the PIP domain compromises pol Eta’s recruitment to sites of DNA damage and thus its ability to resolve the issue. In the UBZ mutant, Pol Eta is likely recruited to the site of damage (e.g. CFS regions) facilitating repair pathway choice but is unable to recruit the necessary (e.g. FANCD2) proteins to alleviate the replication pausing.

## Methods

### Cell Culture

XPPHBE (Coriell cell repository number GM02449) Epstein-Barr Virus transformed XPV lymphoblast cell line (XPV-L Pol eta^−/−^) was grown in RPMI media supplemented with 15% FBS and 1% penicillin and streptomycin. XP30RO SV40 transformed XPV fibroblasts cell line (XPV-F Pol eta^−/−^) was grown in DMEM/F12 media (Gibco) supplemented with 15% FBS and 1% penicillin and streptomycin. XPV fibroblasts stably complemented with WT Pol eta (XPV-F + Pol eta) were generated by complementing XPV-F Pol eta^−/−^ cell line with the Pol eta cDNA [pcDNA 3.1 zeo(-)] and selected in 100 µg/mL Zeocin^44^. No detectable Pol eta protein expression was observed in the XPV-F Pol eta^−/−^ cell line but significantly high level of Pol eta protein expression was detected in the XPV-F + Pol eta cell line^44^. XPV-F Pol eta^−/−^ and XPV-F + Pol eta cell lines were kindly provided by Kristin A. Eckert, Pennsylvania State University College of Medicine, Hershey, PA, USA.

### Single Molecule Analysis of Replicated DNA (SMARD)

SMARD analysis was carried out using a previously described procedure^3,45,46^. Briefly, exponentially growing cells were cultured in media containing 30 μM 5-iodo-2′-deoxyuridine (IdU) at 37°C for 4h (Sigma-Aldrich, St. Louis, MO). After 4h, the cells were centrifuged at 800rpm for 5 min and the media containing IdU was removed. The cells were then cultured in fresh RPMI medium containing 30 μM 5-chloro-2′-deoxyuridine (CIdU) (Sigma-Aldrich, St. Louis, MO) and incubated for an additional 4h. After 4h, the cells were collected by centrifugation and were resuspended at 3 × 107 cells per ml in PBS. The cells were resuspended in an equal volume of molten 1% InCert agarose (Lonza Rockland, Inc., Rockland, ME) in PBS. DNA gel plugs were made by pipetting the cell-agarose mixture into a chilled plastic mold with 0.5- x 0.2-cm wells with a depth of 0.9 cm. The gel plugs were allowed to solidify on ice for 30 min. The cells in the plugs were lysed in buffer containing 1% n-lauroylsarcosine (Sigma-Aldrich), 0.5 M EDTA, and 20 mg/ml proteinase K. The gel plugs were incubated at 50°C for 3 days and treated with fresh proteinase K at 20 mg/ml concentration (Roche Diagnostics), every 24 h.

The Proteinase K digested plugs were then rinsed in Tris-EDTA (TE) and subjected to phenylmethanesulfonyl fluoride (PMSF) (Sigma-Aldrich) treatment. To prepare the cells for restriction enzyme digestion, the plugs were washed with 10 mM MgCl2 and 10 mM Tris-HCl (pH 8.0) and the genomic DNA in the gel plugs was digested with 70 units of PmeI or SbfI (New England BioLabs Inc.) at 37°C overnight. The digested gel plugs were rinsed with TE and cast into a 0.7% SeaPlaque GTG agarose gel (Lonza Rockland, Inc.) for size separation of DNA by pulse field gel electrophoresis. Gel slices from the appropriate positions in the pulsed-field electrophoresis gel were melted at 72°C for 20 min. The melted agarose was digested with GELase enzyme (Epicentre Biotechnologies 1 unit per 50 μl of agarose suspension) by incubating the GELase-DNA-agarose mixture at 45°C for 4 h. The resulting DNA was pipetted along one side of a coverslip that had been placed on top of a 3-aminopropyltriethoxysilane (Sigma-Aldrich)-coated glass slide and allowed to enter by capillary action. The DNA was denatured with sodium hydroxide in ethanol and fixed with glutaraldehyde.

The slides containing the DNA were hybridized overnight with biotinylated probes (represented as blue bars on the CFS locus maps). The next day, the slides were rinsed in 2 × SSC (1× SSC is 0.15 M NaCl plus 0.015 M sodium citrate) 1% SDS and washed in 40% formamide solution containing 2 × SSC at 45°C for 5 min and rinsed in 2 × SSC-0.1% IGEPAL CA-630. Following several detergent rinses (4 times in 4× SSC-0.1% IGEPAL CA-630), the slides were blocked with 1% BSA for at least 20 min and treated with Avidin Alexa Fluor 350 (Invitrogen Molecular Probes) for 20 minutes. The slides were rinsed with PBS containing 0.03% IGEPAL CA-630, treated with biotinylated anti-avidin D (Vector Laboratories) for 20 min, and rinsed again. The slides were then treated with Avidin Alexa Fluor 350 for 20 min and rinsed again, as in the previous step. The slides were incubated with the IdU antibody, a mouse anti-bromodeoxyuridine (Becton Dickinson Immunocytometry Systems), the antibody specific for CldU, a monoclonal rat anti-bromodeoxyuridine (anti-BrdU) (Abcam) and biotinylated anti-avidin D for 1 h. This was followed by incubation with Avidin Alexa Fluor 350 and secondary antibodies, Alexa Fluor 568 goat anti-mouse IgG (H+L) (Invitrogen Molecular Probes), and Alexa Fluor 488 goat anti-rat IgG (H+L) (Invitrogen Molecular Probes) for 1 hour. The coverslips were mounted with ProLong gold antifade reagent (Invitrogen) after a final PBS/CA630 rinse. Fluorescence microscopy was carried out using a Zeiss fluorescence microscope to monitor the IdU/CIdU nucleoside.

### DNA fiber analysis

DNA fibers were stretched and prepared using a modification of a procedure described previously^47,48^. Briefly, cells were pulse labeled with 30 µM IdU/CIdU for 20 minutes each. Labelled cells were resuspended in cold PBS at 1 × 106 cells/ml. Two µl of cell suspension was spotted onto a clean glass slide and to lyse it, 10 μl of spreading buffer (0.5% SDS, 200 mM Tris-HCl (pH 7.4), and 50 mM EDTA) was added. The cells were incubated for 6 minutes, and the slides were tilted to spread the DNA. Slides were either fixed in methanol and acetic acid (3:1) for 2 min, followed by denaturation with 2.5 M HCl for 30 min at room temperature. Alternatively, the DNA was denatured with sodium hydroxide in ethanol and then fixed with glutaraldehyde. The slides were blocked with 1% BSA for at least 20 min. The slides were incubated with the antibodies described above for SMARD. The coverslips were mounted with ProLong gold antifade reagent (Invitrogen) after a final PBS/CA630 rinse. Fluorescence microscopy was carried out using a Zeiss microscope to monitor the IdU/CIdU nucleoside incorporation.

### Comet Assay

Cells were collected by centrifugation at 1000rpm for 5 minutes. Samples were then washed once with 1X PBS and centrifuged again at 1000rpm for 3 minutes. Samples were then resuspended in 1mL of 1X PBS at 1X10^6^ cells/mL. On slides coated with 1% normal melting point agarose, a mixture of 1:7 cell suspension to 1% low melting point agarose was added to each slide and covered with a coverslip. Slides were placed in 4°C for thirty minutes. Cover slips were gently removed, and cells were placed in lysis buffer (5M NaCl, 0.5M EDTA, 10mM Tris-HCl ph10, 1% Triton) for 90 minutes at 4°C. Lysis buffer was removed and denaturation buffer (300mM NaOH, 1mM EDTA) was added for 30 minutes at 4°C. Denaturation buffer was removed, and slides were placed into an electrophoresis unit containing 1X TBE buffer. Electrophoresis was run at 3V/cm for 30 minutes at room temperature. Slides were removed from the electrophoresis unit and put into 2 washes with distilled water for 5 minutes each. Slides were then dehydrated using washes with ice cold 70% ethanol 3 times for 5 minutes each. Slides were left to dry at room temperature and stained with DAPI. Images were acquired using 20X magnification on Zeiss Axio microscope. Data was analyzed using Adobe photoshop tools.

### Immunofluorescence

Cells were collected and seeded on coverslips in 6 well-culture plates at 250,000 cells per coverslip/well in 3ml of culture media. Cells were then treated with 0.4uM Aphidicolin overnight(16h) to induce replication stress. Treatment was withdrawn and media was removed. Coverslips seeded with cells were fixed and permeabilized for 15 minutes with 0.5% Triton in DPBS containing 4% formaldehyde. The coverslips were then washed twice with DPBS-to be used for IF, or they can be stored at 4°C for maximum two weeks.

Primary Antibodies used-Primary antibodies used were anti-FANCD2 (1:1000, NB100-182, Novus), anti-cGAS (1:1000 D1D3G, Rabbit mAb), anti-Phospho-Histone H2A.X (Ser139), anti-XPV DNA Pol eta (Novus NBP1-87165, 1:2000), anti-BRCA1 (Sigma 07-434, 1:1000), anti-BRCA2 (Merck Millipore OP95, 1:1000) and anti-RAD51 (CosmobioUSA BAM-70-002-EX, 1:1000).

### Western Blotting

Cells were harvested after treatment by centrifugation followed by PBS wash. Cells were lysed with 2X Laemmli buffer and lysates were denatured at 100°C for 15 minutes. Proteins were separated using NuPAGE 4-12% Bis-Tris Mini Gels (Thermo Scientific) or 3-8% Tris-Acetate mini gels (Thermo Scientific) and transferred to nitrocellulose membrane. Membranes were blocked with AdvanBlock (Advansta) blocking buffer for 1 hour at room temperature and then incubated with primary antibodies overnight at 4°C. Membranes were washed 3 times for 15 minutes and then incubated with horseradish peroxidase (HRP)-linked secondary antibodies for 1 hour at room temperature. Proteins were then detected by chemiluminescence.

Time Course Experiment: Cells were grown in corresponding media and treated with 2mM HU for 4 hours. Cells were released from HU into fresh media and collected at various time points: Immediately after release (time 0, 4, 8, 12,24, 32, and 48 hours after release. Cells were then harvested and lysed for immunoblotting. Primary Antibodies for western blot included anti-DNA Polymerase Eta (Cell Signaling-13848S, 1:2000), anti-yH2AX (Cell Signaling – 9718T), anti-pCHK1 (s317) (Cell signaling –123402), anti-H3 (Cell Signaling – 12648T), anti-PCNA mAB (Cell Signaling-2586T, 1:1000), anti-ubiquityl PCNA mAB (Cell Signaling-13439S, 1:1000), anti-FANCD2 (SantaCruz, 1:2500), phospho-ATR (Cell Signaling, 30632, 1:2000)

### Non-Denaturing ssDNA gap detection by immunofluorescence

Cells were grown in the presence of 30uM IdU for 48 hours before treatment with 2mM hydroxyurea. Cells were then collected by centrifugation and washed with PBS (Phosphate Buffered Saline). Cells were seeded in glass slides by cytospin for 6 minutes at 1100 RPM. Cells were fixed and permeabilized with 0.5% Triton X-100 and 4% paraformaldehyde in PBS at room temperature for 15 minutes. Fixed cells were then blocked with 3% BSA for 1 hour at room temperature. Cells were incubated with primary antibodies against pRPA at 4°C overnight. Cells were then washed 3 times with PBS at room temperature followed by incubation with antibodies against IdU for 1 hour at room temperature. Cells were washed 3 times with PBS followed by secondary antibodies for 1 hour at room temperature. Cells were then mounted with Prolong with DAPI.

### Mitotic DNA Synthesis (MiDAS)

Harvesting mitotic cells: To augment mitotic cells, asynchronous cells were treated with CDK1 inhibitor RO-3306 (Merck) at a final concentration 7 μM for 7:30 h, or in the last 7:30h of APH treatment as and when needed. The cells were then washed with pre-warmed Edu-containing medium (37 °C) for three times within 5 minutes before being released into pre-warmed fresh medium with Edu at a final concentration of 20 μM for 45mins-1 h. This allows the cells to advance into mitosis. The mitotic population is then analyzed. Next, the cells are collected and centrifuged at RT at 350g/rcf for 5 minutes to form a pellet. They are then washed in pre-warmed fresh medium two times before resuspending them into DPBS (1 million cells/ml of DPBS). Further, cells are seeded on Corning-single frosted microslides using Cytospin 4 (Thermoscientific) for 6 minutes at 1100rpm at medium acceleration. Once cells are seeded, they are fixed and permeabilized for 15 minutes with 0.5% Triton in DPBS containing 4% formaldehyde. The slides are then washed twice with DPBS-to be used for EdU labelling and IF, or they can be stored at 4 °C for a maximum of two weeks.

EdU detection and IF staining for MIDAS: The cells were blocked using blocking buffer (3% BSA in 1X PBS) for 2 hours at RT. To detect EdU Click-IT chemistry was used as per the manufacturer’s instructions (Click-iT™ EdU Cell Proliferation Kit for Imaging, Alexa Fluor™ 594 dye). EdU detection was coupled with IF staining. The cells were incubated with primary antibody diluted in the blocking buffer at 4 °C overnight; followed by three washes 15 mins each with DPBS. Next, the cells were incubated with secondary antibodies diluted in the blocking buffer for 1 hour at RT, followed by three washes, 15 mins each with DPBS. Slides were then mounted with Prolong gold containing DAPI (Invitrogen Prolong Gold anti fade reagent with DAPI, P36935). The slides were allowed to dry overnight prior to IF imaging. Images were captured using-63mm oil immersion lens, Ziess Axio microscope. Primary and Secondary antibodies for MiDAS experiments: Primary antibodies used were FANCD2 (1:1000, NB100-182, Novus). Secondary antibodies used were Anti-rabbit IgG Fab2 Alexa Fluor ®594 (1:20000, 8889S, Molecular probes).

#### Analysis

Under-replicated DNA at CFS: Mitotic cells were labelled for EdU and IF staining was performed. FANCD2 twin foci were used as a marker of CFS location. Each mitotic cell was analyzed for EdU labelled FANCD2 twin foci. Each FANCD2 twin foci showed 1-2 EdU foci suggesting DNA replication at under-replicated regions called CFS.

#### Cytoplasmic dsDNA Analysis

Chromatin fragment staining positive for DAPI, proximal to parent nucleus was analyzed as cytoplasmic DNA. These fragments were stained for pico green (Pico488 dsDNA Quantification Reagent, Lumiprobe-42010) that mark damaged regions as well as for sensing cytosolic dsDNA. This was used to assess the fate of post-mitotic consequesnces to determine persistence of genome instabilty in different pol eta mutant cell lines.

### Quantification and Statistical Analysis

All statistical calculations were performed using GraphPad Prism v.9.0. A two-sided Student’s t-test was used to calculate p-values in GraphPad Prism v.9.0. All the experiments were generally conducted with at least three biological replicates. Differences were considered significant at p-values: * <0.03, ** <0.0021, *** <0.0002, **** <0.0001.

## Author Contributions

The project was conceived by A.M. Most of the experiments were conducted by M.N, A.B.G and V.K. Data analysis was carried out by M.N, A.B.G, J.E.G, V.K, A.P and A.J. The data was analyzed and discussed by A.M, M.N and J.G.

## Acknowledgements

We thank Kristin A. Eckert at Pennsylvania State University College of Medicine for XPV-F Pol eta^−/−^ and XPV-F + Pol eta fibroblasts analyzed as part of this study. This work was supported by the National Institutes of Health (NIH) grants 3R00HL136870-04S1 and 5R00HL136870-04 to A.M.

## Author Information

The authors have NO commercial affiliations or conflicts of interest.

## Notes

### Competing Interest Statement

The authors have declared no competing interest.

